# Regional diversity in the postsynaptic proteome of the mouse brain

**DOI:** 10.1101/368910

**Authors:** Marcia Roy, Oksana Sorokina, Colin McLean, Silvia Tapia-González, Javier DeFelipe, J. Douglas Armstrong, Seth G. N. Grant

**Author notes:** Equal contribution (Marcia Roy, Oksana Sorokina).

## Abstract

The proteome of the postsynaptic terminal of excitatory synapses comprises over one thousand proteins in vertebrate species and plays a central role in behavior and brain disease. The brain is organized into anatomically distinct regions and whether the synapse proteome differs across these regions is poorly understood. Postsynaptic proteomes were isolated from seven forebrain and hindbrain regions in mice and their composition determined using proteomic mass spectrometry. Seventy-four percent of proteins showed differential expression and each region displayed a unique compositional signature. These signatures correlated with the anatomical divisions of the brain and their embryological origins. Biochemical pathways controlling plasticity and disease, protein interaction networks and individual proteins involved with cognition all showed differential regional expression. Combining proteomic and connectomic data shows that interconnected regions have specific proteome signatures. Diversity in synapse proteome composition is key feature of mouse and human brain structure.

## Introduction

Synapses are the specialized junctions between nerve cells and are present in vast numbers in the mammalian nervous system. During the 1990s, synapses were thought to be relatively simple connectors, but the application of proteomic mass spectrometry in 2000 revealed an unanticipated complexity in their protein composition^1^. Both the presynaptic and postsynaptic proteomes have since been systematically characterized in several vertebrate species and thousands of proteins have been identified^2-13^. Phosphoproteomic studies have shown that neural activity causes changes in large numbers of proteins^14,15^. These findings have transitioned the view of synapses to one where they are highly sophisticated and complex signaling machines that process information. The importance of understanding this complexity is underscored by the finding that over 130 human brain diseases are caused by mutations disrupting postsynaptic proteins^16,17^.

It is of fundamental importance to understand how the high number of postsynaptic proteins are organized physically (within synapses) and spatially (between synapses). Biochemical studies have shown that postsynaptic proteins are typically assembled into a hierarchy of complexes and supercomplexes (complexes of complexes)^18-20^. The prototype of postsynaptic supercomplexes are those formed by the scaffolding protein PSD95 (also known as Dlg4). Dimers of PSD95 assemble with complexes of neurotransmitter receptors, ion channels, signaling and structural proteins into a family of high molecular weight (1-3 MDa) structures in excitatory synapses^18-20^. PSD93 (also known as Dlg2), which is a paralog of PSD95, co-assembles with PSD95 to bind NMDA receptors and these are an important functional subset of the PSD95 supercomplex family. Other combinations of proteins form other subtypes of PSD95 supercomplexes, such as those containing potassium channels and serotonin receptors^20-23^. Together, these members of the PSD95 supercomplex family confer diverse signal processing functions to the synapse.

The principles underlying the spatial organization of synapse proteomes in the brain is less well understood. To date, most studies of the synapse proteome have focused on defining composition from limited regions of the brain (or the whole brain). However, at the macroscopic level, brain architecture is characterized by regions with distinct functions^24^. It is therefore of importance to ask if synapse proteomes differ between brain regions and whether any differences might be relevant to their function or to the connectivity between these regions. In a recent study, we reported that regions of human neocortex differ in the composition of their postsynaptic proteomes and that these compositional differences correlate with functional properties^25^. The present study employs a similar analysis applied to the mouse brain, which allows us to ask if conserved principles may apply across these two species that evolved from a common ancestor ~90 million years ago.

Using a method suitable for the isolation and direct quantification of mouse synapse proteomes from small amounts of brain tissue, we compare and contrast the synapse proteomes isolated from seven integral regions of the adult mouse brain. The postsynaptic proteome was analyzed to a depth of 1,173 proteins and differential expression signatures were identified and characterized in each brain region. We use these datasets to analyze the spatial organization of the postsynaptic proteome in the mouse brain and identify organizational principles shared with humans. This large-scale dataset is a useful resource for the field of neuroscience and future studies using mouse models of human disease.

## Materials and Methods

### Dissections of mouse brain regions

This study was performed using 8-week-old male C57BL/6J mice. All experimental protocols involving the use of animals were performed in accordance with recommendations for the proper care and use of laboratory animals and under the authorization of the regulations and policies governing the care and use of laboratory animals (EU directive n° 86/609 and Council of Europe Convention ETS123, EU decree 2001-486 and Statement of Compliance with Standards for Use of Laboratory Animals by Foreign Institutions n° A5388-01, National Institutes of Health, USA).

The mice (*n*=6) were anesthetized with a pentobarbital dose of 40 mg/kg body weight and sacrificed by decapitation. The brains were rapidly removed and kept on ice while the areas of interest were dissected from the right hemisphere using the microdissection method of Palkovits^37^. Large regions were collected from the frontal, medial and caudal cortex, as well as the right caudate putamen, right hippocampus, whole hypothalamus, and cerebellum (right half), which was cut previously through the vermis (Supplementary Figure 1). The samples were frozen on liquid nitrogen and stored at −80°C until processed.

### PSD isolation and protein preparations for mass spectrometry

Dissected mouse brain regions were homogenized by performing 12 strokes with a Dounce homogenizer containing 2 mL ice-cold homogenization buffer (320 mM sucrose, 1 mM HEPES, pH 7.4) containing 1x Complete EDTA-free protease inhibitor (Roche) and 1x Phosphatase inhibitor cocktail set II (Calbiochem). Synaptosomes were isolated from homogenized mouse brain tissue as described^2^. Briefly, insoluble material was pelleted by centrifugation at 1,000 x g for 10 minutes at 4°C. The supernatant (S1) was removed and the pellet resuspended in 1 mL homogenization buffer and an additional six strokes were performed. Following a second centrifugation at 1,000 x g for 10 minutes at 4°C, the supernatant (S2) was removed and pooled with S1. The combined supernatants were then centrifuged at 18,500 x g for 15 minutes at 4°C. The pellet was resuspended in 0.25 mL homogenization buffer and 0.25 mL extraction buffer (50 mM NaCl, 1% DOC, 25 mM Tris-HCl, pH 8.0) containing 1x Complete EDTA-free protease inhibitor (Roche) and 1x Phosphatase inhibitor cocktail set II (Calbiochem) and incubated on ice for 1 hour. The resulting PSD extracts were centrifuged at 10,000 x g for 20 minutes at 4°C and the resulting supernatant filtered through a 0.2 μm syringe filter (Millipore).

### Sample preparation and LC-MS/MS analysis

All chemicals were purchased from Sigma-Aldrich unless otherwise stated. Acetonitrile and water for HPLC-MS/MS and sample preparation were HPLC quality and were purchased from Thermo Fisher. Formic acid was supra-pure (90-100%) purchased from Merck while trypsin sequencing grade was purchased from Promega. All HPLC-MS connector fittings were either purchased from Upchurch Scientific (Hichrom) or Valco (RESTEK). Proteins were acetone precipitated, protein pellets reconstituted in SDS-PAGE loading buffer, and briefly run on a 4-12% Bis-Tris gradient gel (Invitrogen) for ~10 minutes. Proteins were in-gel digested using a method similar to that of Shevchenko et al. (2006)^38^. Resulting peptide extracts were then acidified with 7 μL 0.05% TFA and were filtered with a Millex filter (Millipore) before HPLC-MS analysis. Nano-HPLC-MS/MS analysis was performed using an on-line system consisting of a nano-pump (Dionex Ultimate 3000, Thermo Fisher) coupled to a QExactive instrument (Thermo Fisher) with a pre-column of 300 μm x 5 mm (Acclaim Pepmap, 5 μm particle size) connected to a column of 75 μm x 50 cm (Acclaim Pepmap, 3 μm particle size). Samples were analyzed on a 90-minute gradient in data-dependent analysis (one survey scan at 70k resolution followed by the top ten MS/MS).

### Mass spectrometry and data analysis

Data from MS/MS spectra were searched using MASCOT version 2.4 (Matrix Science Ltd) against the *Mus musculus* subset of the National Center for Biotechnology Information (NCBI) protein database (382,487 protein sequences) with maximum missed-cut value set to 2. The following features were used in all searches: i) variable methionine oxidation, ii) fixed cysteine carbamidomethylation, iii) precursor mass tolerance of 10 ppm, iv) MS/MS tolerance of 0.05 amu, v) significance threshold (*p*) below 0.05 (MudPIT scoring) and vi) final peptide score of 20.

Progenesis version 4 (Nonlinear Dynamics, UK) was used for HPLC-MS label-free quantitation. Only MS/MS peaks with a charge of 2+, 3+ or 4+ were taken into account for the total number of ‘Feature’ (signal at one particular retention time and m/z) and only the five most intense spectra per ‘Feature’ were included. Each LC-MS run is normalized by multiplying a scalar factor. The scalar factor is a ratio in log space of the median intensity of the selected features against the median intensity of the selected feature of a reference spectrum. The associated unique peptide ion intensities for a specific protein were then summed to generate an abundance value and transformed using an ArcSinH function. Based on the abundance values, within group means were calculated and from there the fold changes (in comparison to control) were evaluated. One-way analysis of variance (ANOVA) was used to calculate the *p*-value based on the transformed abundance values. *P*-values were adjusted for multiple comparisons and were calculated either from Progenesis version 4 (Nonlinear Dynamics) or using R (R Core Team, 2013)^39^ based on Benjamini and Hochberg (1995)^40^. Further analysis was performed by extracting a Z-score calculated on ArcSinH average group.

Differentially expressed proteins were only considered significant in the current study if the following conditions were fulfilled: i) adjusted *p*-values (pairwise) less than 0.05, ii) number of unique peptides detected and used in quantification per protein was at least 2 for the 1,173 dataset, and iii) absolute fold change was at least 1.3 for differentially abundant proteins and ≤ 0.667 for downregulated proteins.

### Bioinformatic analyses

The majority of the analysis was performed in R. PCA and Tukey test was performed with R package *FactoMineR* and correlation analysis with the package *corrplot*. Differential stability (DS) analysis was performed as described^41^; briefly, for each protein from the list of 1,173 the average Pearson correlation coefficient was estimated from 14 pairwise Pearson coefficients for six brain samples. Heatmaps were generated with use of the *heatmap.2* function from *gplot* R library: parameters were set to default values with the exception of label and dendrogram visualization control. Hierarchical clustering validation and comparison of dendrograms were performed with package *dendextend*^42^. The number of stable clusters was independently assessed with *nbclust* package^43^, which provides the list of indices to determine the optimal number of clusters. We selected a set of six clusters [the postsynaptic proteome modules (PPMs)] based on the best combination of indices provided by *nbclust* R package. Individual proteins in each of the six PPMs and their abundances across all seven integral regions are listed in Supplementary Table 5 and proteins in each module were ranked by their abundance in each of the seven regions of the mouse brain examined.

We used Bioconductor package ClusterProfiler for Gene Ontology (GO) and KEGG enrichment analysis^44^ and Bioconductor ReactomePA package for pathway over-representation analysis (http://bioconductor.org/packages/release/bioc/html/ReactomePA.html).

GO function and KEGG pathway enrichment for all proteins was performed using DAVID (https://david.ncifcrf.gov/). Disease enrichment for each brain region and each protein module was performed using DAVID (https://david.ncifcrf.gov/) and KEGG pathway enrichment was then performed by searching ranked protein lists obtained using GSea version 2.1.0 (http://software.broadinstitute.org/gsea/index.jsp) as previously described^45^.

Circular hierarchical clustering of protein modules for the visualization of inter-region molecular interactions was performed using Circos (http://circos.ca)^46^. A Circos configuration file was created representing brain regions as “karyotypes”. All proteins were grouped into “modules” according to their abundance similarity. Proteins that have positive abundance in more than one region were shown as links between regions. The width of each link is proportional to the fraction of the regional proteins that contributed to the link, while its color corresponds to that of the respective “module”. All preprocessing of the relative abundance information and generation of appropriate Circos files were performed in R. Scripts are available on request.

For DS analysis, we used the described approach^41^ on the MS intensity values obtained for all 1,173 proteins identified with a minimum of two unique peptides in order to identify synaptic proteins with highly reproducible expression patterns across all six independent mouse brains. The average pairwise Pearson correlation *ρ* over the six individual mouse brains was quantified and obtained DS values ranged from 0.98 to −0.89.

For correlation with mesoscale mouse connectome data^29^, the mean voxel sum for each region was calculated with respect to all other regions and itself. The correlation of this matrix was then estimated with the matrix of protein abundances. The results are listed in Supplementary Table 3.

### Protein interaction identification and mapping

The full postsynaptic proteome network was built from the list of 1,173 proteins obtained in this study and PPIs obtained by mining publicly available databases: BioGRID^32^, IntAct^33^ and DIP^34^ both for mouse and human. The total network consists of 1,016 proteins and 8,105 PPIs. We applied weights to each interaction based on abundance values for specific brain regions as follows: mean (ExpA, ExpB), so that for each of the regions the specific weight for each of the interactions could be determined. Having varying abundances for interacting proteins in different brain regions, we estimated the region-specific edge that resulted in region-specific PPI. Each brain region network was clustered making use of the spectral properties of the network; the network being expressed in terms of its eigenvectors and eigenvalues, and partitioned recursively (using a fine-tuning step) into communities based on maximizing the clustering measure modularity^47-49^; the modularity of the networks was found to be 0.28-0.42. Enrichment for biological process and cellular component was performed using the topGO package (https://bioconductor.org/packages/release/bioc/html/topGO.html), while functional enrichment of synaptic proteins/gene groups that are known risk factors for schizophrenia was performed using the published schizophrenia risk factor dataset^50^.

The stability of all interactions across the region were assessed by comparison of clustering results for each of seven region-specific networks and assigning each interaction a score of 0 if both proteins appeared in the same cluster and a score of 1 if it appeared in a different cluster. Scores were summed over all seven regions, resulting in sums ranging from 0 (proteins remain in the same cluster in all regions) to 7 (proteins never appear in the same cluster). For the “stable” network, we selected interactions with scores ≤ 2, which means that they persist in the same cluster in 5/7 (70%) regional networks.

PPI networks were visualized with Gephi (https://gephi.org).

For disease enrichment analysis, the community and protein robustness values within the range 0-1 were taken as edge weights. Each region network was then clustered and cluster enrichment was assessed using the TopOntop package (https://github.com/hxin/topOnto) and OMIM/Ensemble Var/genetic annotation data. For disease enrichment the annotation data were standardized using MetaMap^51-53^ and NCBO Annotator (https://www.bioontology.org/annotator-service) to recognize terms found in the Human Disease Ontology (HDO)^54^. Recognized enriched disease ontology terms were then associated with gene identifiers and stored locally. Disease term enrichment could then be calculated using the topology-based elimination Fisher method^55^ found in the topGO package (htt://topgo.bioinf-mpi-inf.mpg.de/), together with the standardized OMIM/GeneRIF/Ensembl variation gene-disease annotation data (17,731 gene-disease associations), and the full HDO tree (3,140 terms). Each region was then examined individually by performing the clustering analysis and enrichment for each of the clusters identified in each of the seven brain regions.

## Results

### Quantification of postsynaptic proteins from brain regions

Seven integral brain regions within the forebrain (prosencephalon) and hindbrain (rhombencephalon) were dissected from six 8-week-old C57BL/6J mice (Figure 1A, Supplementary Figure 1). Forebrain regions included telencephalic structures: frontal cortex (CxF), medial cortex (CxM), caudal cortex (CxCA), hippocampus (Hip), striatum (ST); the hypothalamus (Hyp) represented a diencephalic structure and the hindbrain was represented by cerebellum (CB). PSD fractions were prepared from six mice and all 42 samples were analyzed using LC-MS/MS. Label-free quantitation of peptide intensity identified 1,173 proteins across all seven brain regions (Supplementary Table 1). We found a significant overlap between our dataset and those obtained in other mouse studies^4-6,10-12,22^ (Supplementary Figure 2A,B); the 61 proteins unique to this study are summarized in Supplementary Table 2.

**Figure 1.**
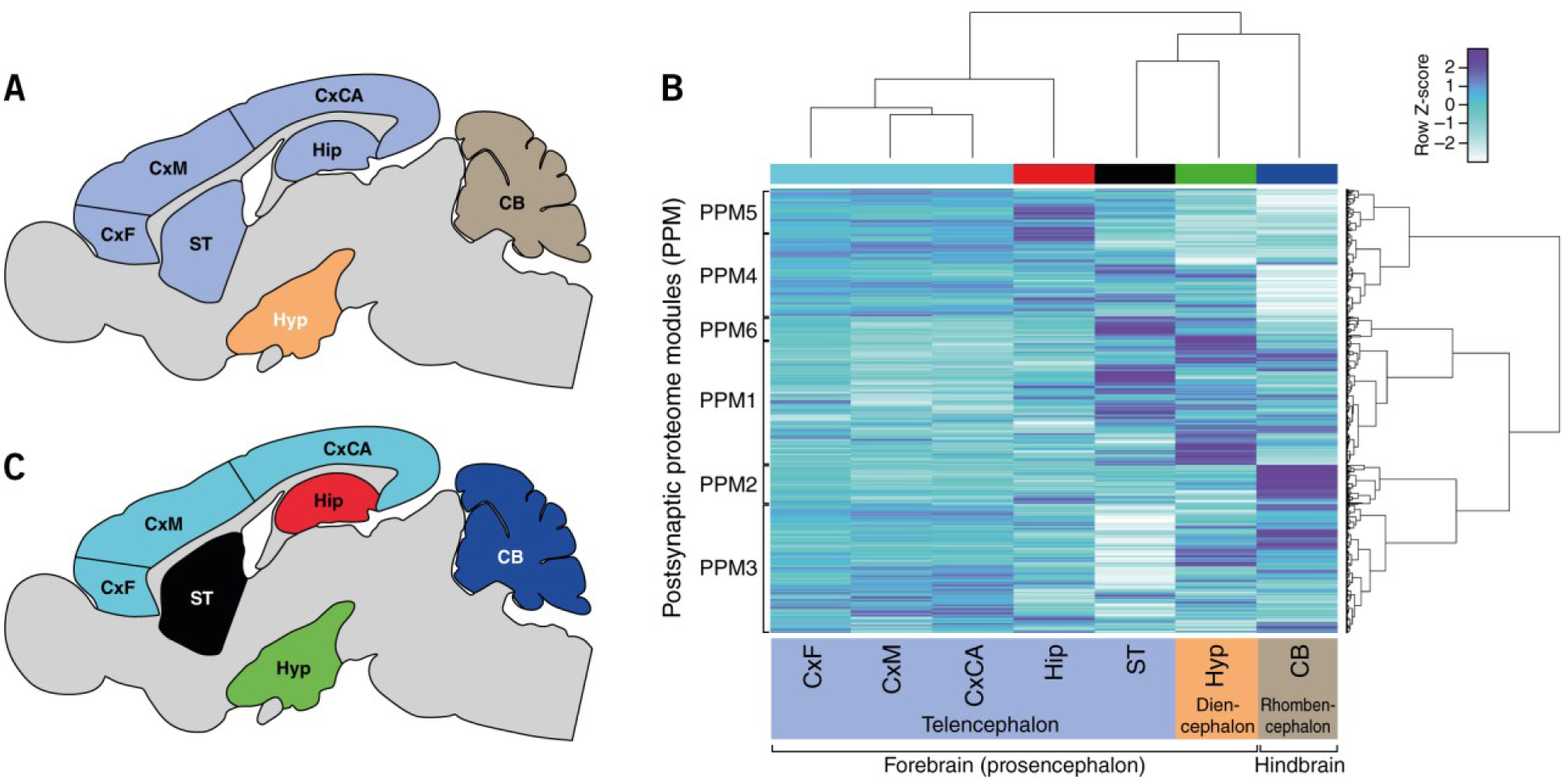
Signatures of postsynaptic proteome composition in mouse brain regions. (A) Seven integral brain regions in mouse: frontal cortex (CxF), medial cortex (CxM), caudal cortex (CxCA), hippocampus (Hip), striatum (ST), hypothalamus (Hyp), cerebellum (CB). Color coded according to vertebrate embryological regions (as in B). (B) Hierarchical clustering by region (*x*-axis) and protein abundance (*y*-axis) shows that each region has a unique signature of postsynaptic proteome composition. (C) Neuroanatomical map of clusters in (B) showing proteome organization into forebrain and hindbrain structures: telencephalon, diencephalon and rhombencephalon.

To determine the validity of pooling data from six mice we performed several analyses. We first used the differential stability (DS) approach, which has been previously applied to transcriptomic and proteomic analyses of adult human brain regions^25,26^ (Supplementary Table 3). For this we estimated the average pairwise Pearson correlation to identify the proteins that demonstrate similar patterns across all six brains. From 1,173 synaptic proteins, roughly half (572) displayed a high DS correlation, or similar expression patterns across all brain regions (Supplementary Table 3A,B). We found that these were functionally enriched in synaptic transmission proteins (*q* = 8.25 x 10^−19^), ATP metabolic processes (*q* = 1.33 x 10^−12^) and calcium ion transporting proteins (*q* = 2.02 x 10^−9^). Proteins involved in pathways associated with learning (*q* = 8.08 x 10^−6^), memory (*q* = 6.23 x 10^−3^) and behavior (*q* = 1.39 x 10^−7^) were also over-represented in this high DS subset. Components of several KEGG pathways were also highly correlated between individuals, including long-term potentiation (*q* = 7.95 x 10^−7^), calcium signaling pathways (*q* = 1.13 x 10^−5^), Huntington’s (*q* = 2.66 x 10^−7^), Alzheimer’s (*q* = 2.14 x 10^−5^) and Parkinson’s disease (*q* = 8.95 x 10^−6^) (Supplementary Table 3C). The most highly conserved proteins between the six individual mice with the highest DS values were STX1A (*p* = 0.96), STUM (*p* = 0.96), CDH13 (*p* = 0.95) and ATP1B2 (*p* = 0.95) (Supplementary Table 3B). We also compared the distribution of synaptic protein abundances across individual mice by principal component analysis (PCA). This analysis (Supplementary Figure 3A) indicates that the synaptic proteome of the mice largely overlap; with brains A-C corresponding to the central region of the distribution. As Tukey’s HSD test shows no significant difference in the mean values of six individuals at a confidence level of 95% (Supplementary Figure 3B), we determined that the data from all six individuals could be combined and the mean protein abundances were then used for all downstream analyses.

### Regional differences in postsynaptic proteome composition

To identify postsynaptic proteins with differential expression between brain regions, proteins having a mean peptide intensity of 1.5-fold or greater in one brain region compared with any other and determined to be significant with *p* < 0.05 were identified (Supplementary Table 4). Eight hundred and sixty-eight (74%) proteins were found to be differentially expressed in at least one region compared with all others (Supplementary Table 4). The regions with the largest number of differentially expressed proteins were the cerebellum (251), hypothalamus (243) and striatum (161). By contrast, the frontal (14), medial (70) and caudal cortex (34) were found to contain the lowest number of differentially expressed proteins compared with all other regions (Figure 1C).

Hierarchical clustering of all proteins revealed that each region has a unique signature of expression. Moreover, these signatures are organized in line with the classical anatomical architecture of the brain: the three cortical regions showed greatest similarity, and the next most similar region was the hippocampus, then striatum, hypothalamus and cerebellum (Figure 1B). This clustering reflects the embryological divisions of the vertebrate brain into telencephalon, diencephalon and rhombencephalon (Figure 1C). Moreover, these results complement findings in the human neocortex, where unique signatures were also found for each region^25,27^. Together, these findings indicate that compositional differences in the postsynaptic proteome reflect, at least in part, the embryological patterning mechanisms that define brain regions.

This clustering approach also allowed us to examine region-specific functions. We identified six sets of proteins, which we call postsynaptic proteome modules (PPM 1-6) (Figure 1B, Supplementary Table 5). As indicated by the clustering and Circos plots (Supplementary Figure 4), these PPMs were differentially distributed in brain regions. To understand the functional significance of differential protein expression in modules and regions, we analyzed the KEGG biochemical and disease pathways in PPMs (Figure 2A). The PPMs showed differential composition of pathways. For example, neurodegenerative diseases were found in PPM1, whereas synaptic plasticity (long-term potentiation, long-term depression) and relevant signaling pathways were in PPM2.

**Figure 2.**
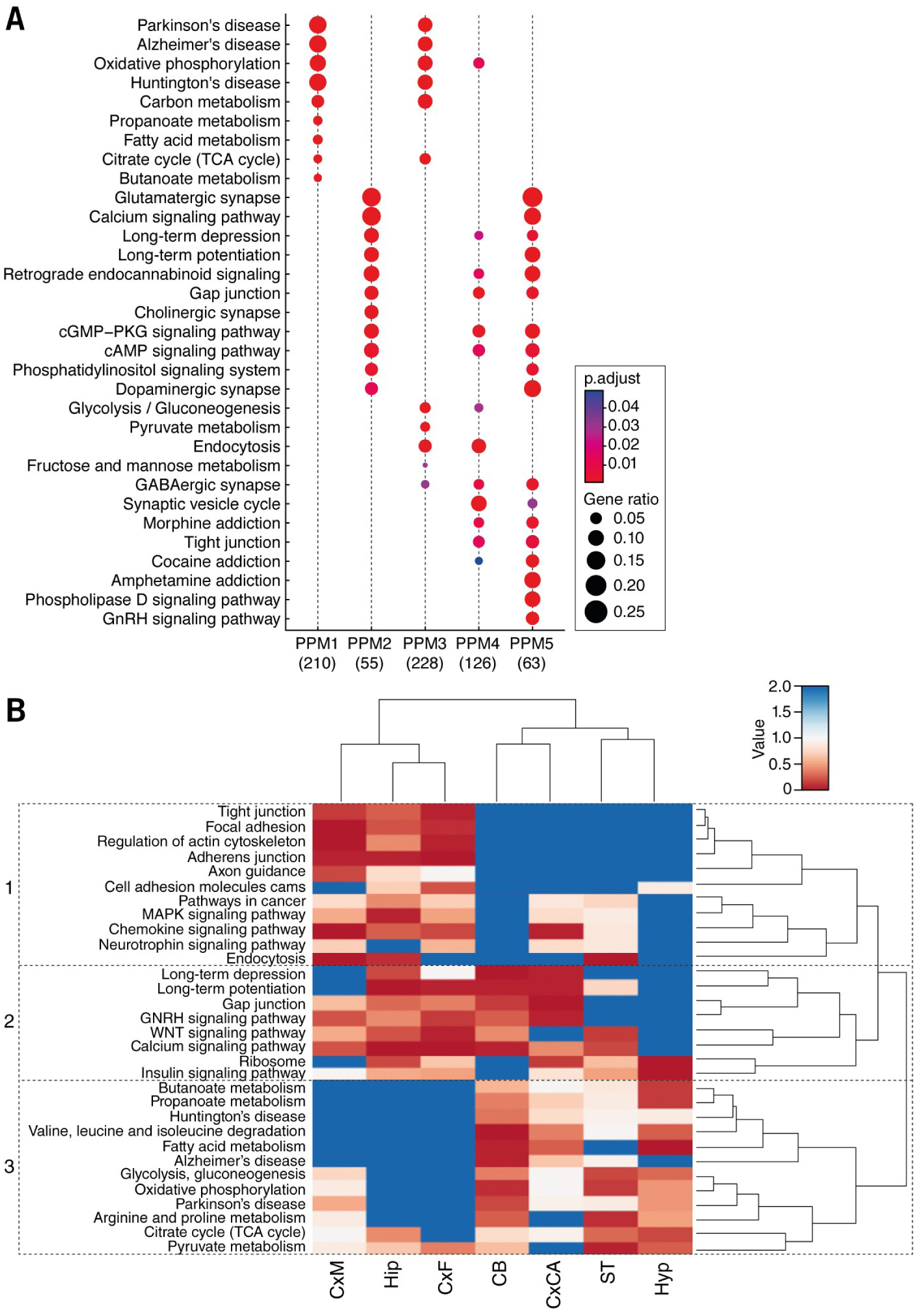
Biochemical pathways and functions in brain regions and postsynaptic proteome modules (PPMs). (A) KEGG pathway term (*y*-axis) enrichment in PPMs (*x*-axis) with the number of proteins contributing to KEGG enrichment indicated in brackets. Size of the dots represents the number of genes associated with that pathway (GeneRatio) and the significance indicated by the p-adjust color bar. (B) Heatmap of the KEGG biochemical pathway and disease enrichment terms (*y*-axis) based on the ranked abundance of postsynaptic proteins in each region (*x*-axis). Three clusters of KEGG terms are boxed: 1) many signal transduction mechanisms, 2) synaptic plasticity and other signaling processes, 3) neurodegenerative diseases and metabolic mechanisms.

Examination of KEGG pathway enrichment in brain regions (Figure 2B) revealed three major groups. It is striking that very similar groupings were observed in the analysis of human neocortical regions^27^. Group 1 contained terms including MAPK, chemokine, neurotrophin pathways; group 2 included synaptic plasticity mechanisms and calcium signaling; group 3 included neurodegenerative diseases (Alzheimer’s, Huntington’s, Parkinson’s) and metabolic mechanisms (glycolysis/gluconeogenesis, oxidative phosphorylation). These findings suggest that biochemical pathways in the postsynaptic proteome are differentially distributed across brain regions and that the mechanisms controlling this distribution are species conserved.

### Distribution of mechanism of cognition and protein complexes

The seven regions of the brain examined in this study are thought to play distinct but interdependent roles in cognitive function. Therefore, we examined the distribution of 33 selected proteins that are known to play roles in cognition (Figure 3A). Hierarchical clustering shows that the three cortical regions examined (CxF, CxM, CxCA) cluster together by similarity, while the Hip region clusters separately from all others. The Hyp and ST regions cluster together by similarity in their abundances of proteins involved in memory and cognition, while the CB clusters separately from all of the other six regions. The abundances of these proteins clustered into two main branches (Figure 3A).

**Figure 3.**
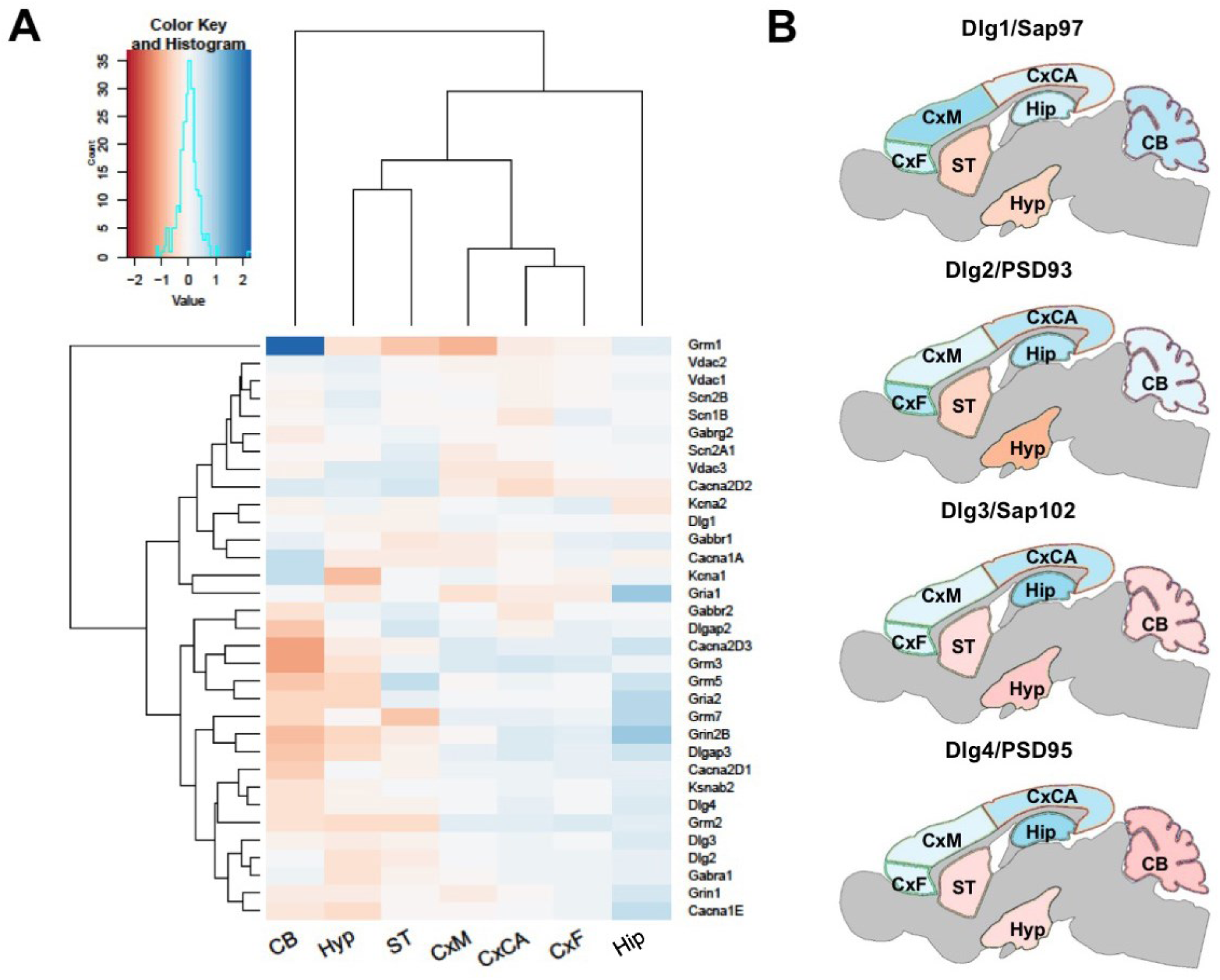
Distribution of 33 selected proteins known to play roles in cognition. (A) Hierarchical clustering indicates that the three cortical regions (CxF, CxM and CxCA) and Hyp and ST regions cluster together by similarity, but separately from Hip CB clusters separately from all of the other six regions. (B) The abundances of the four MAGUK scaffold protein paralogs Dlg1 (Sap97), PSD93 (Dlg2), Dlg3 (Sap102) and PSD95 (Dlg4) mapped across the various brain regions.

In order to assess the heterogeneity of synaptic protein complexes throughout the brain, the abundances of the four MAGUK scaffold protein paralogs Dlg1 (also known as Sap97), PSD93 (Dlg2), Dlg3 (also known as Sap102) and PSD95 (Dlg4) were mapped across the various brain regions. We found that these four molecules, which play fundamental roles in synaptic transmission, were differentially distributed throughout the brain, with Dlg1 being most abundant in the synapses of the CxM and PSD93 most abundant in Hip, CxCA and CxF. By contrast, both Dlg3 and PSD95 showed similar protein abundance profiles across the various brain regions (Fig. 3B).

### Correlations of regional synapse proteomes with the connectome

There are large-scale efforts to map the mouse brain connectome by identifying the projections of neurons between brain regions^28,29^. Because these connections are made at synapses, it follows that there may be a relationship between the molecular composition of synapses in one region and their interconnections. To address this, we asked if the synaptic proteins quantified in this study correlated with connectivity data from the Allen Brain Institute’s Mouse Brain Connectivity Atlas (mesoscale connectome)^29^ (Materials and Methods, Figure 4, Supplementary Table 6). Hierarchical clustering of postsynaptic proteome abundance and connection strength approximated from projection volume shows that regional connections are associated with distinct signatures of proteins. Moreover, two major branches separated cortex, striatum and hippocampus from cerebellum and hypothalamus, suggesting that hindbrain and basal forebrain connections have broadly distinct molecular properties compared with connections of other forebrain structures.

**Figure 4.**
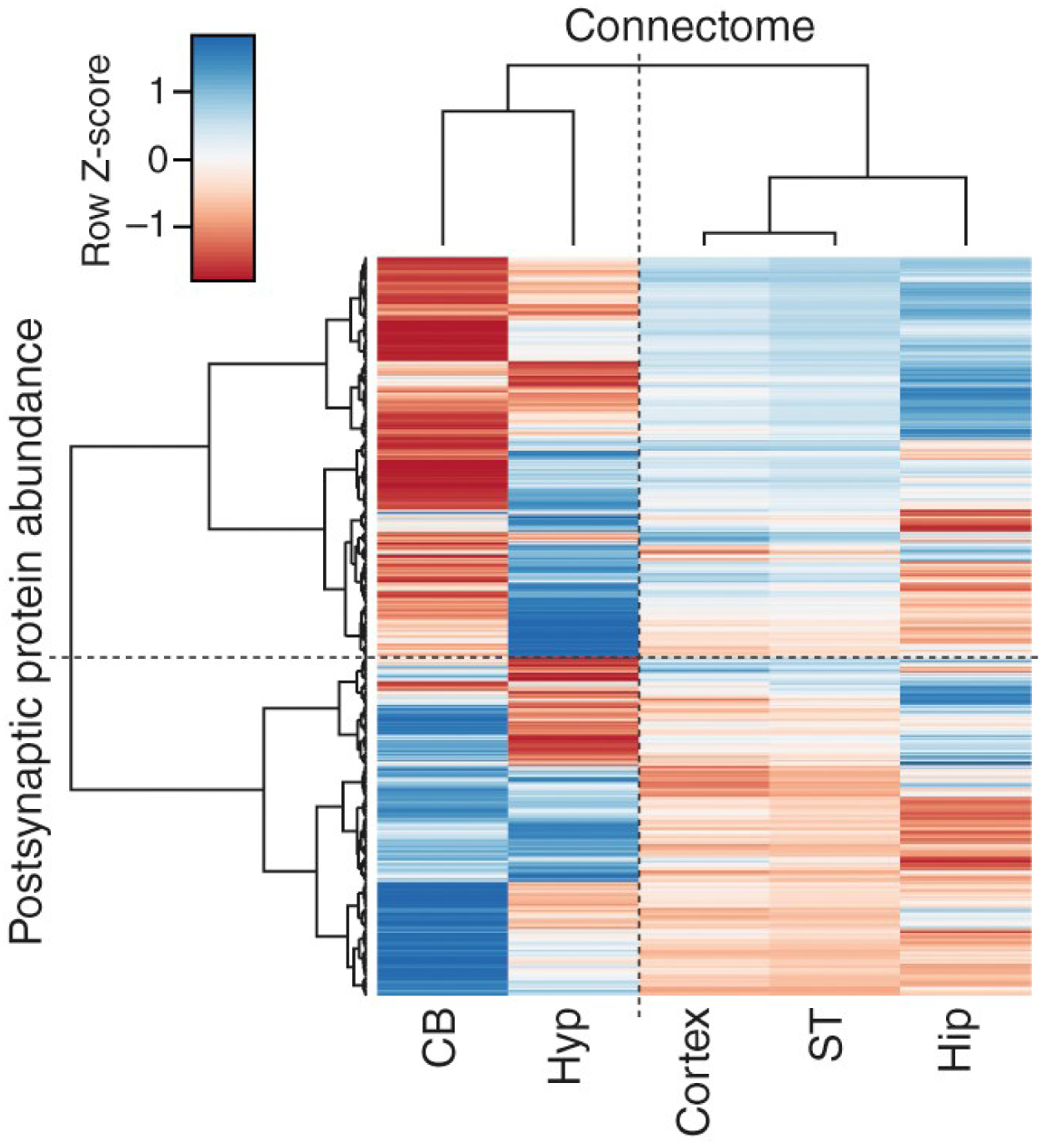
Correlation between brain region-specific postsynaptic protein abundance and mesoscale connectome. Clustering heatmap of the correlation between protein abundance and neuron projection volume in each brain region. Color key shows Z-transformed correlation values; red corresponds to negative correlation and blue to positive correlation.

We found that, in the hippocampus, Dlg3, PSD93 and PSD95 but not Dlg1 were highly correlated (R^2^ = 0.7-0.8) with projection volume (Supplementary Table 6). We then asked if the biochemical pathways that underlie brain connectivity were brain region specific, and performed functional enrichment on the synaptic proteins that were highly correlated (R^2^ ≥ 0.6) with neuron projection volume for each region. Pathways associated with glutamatergic synapses, calcium signaling, long-term potentiation (LTP), long-term depression (LTD) and insulin signaling were over-represented in the hippocampus, striatum and cortical regions, whereas pathways involved in Parkinson’s, Huntington’s and oxidative phosphorylation and mitochondrial components were enriched in the cerebellum and hypothalamus. Additionally, the molecular correlates of connectivity in the hippocampus are uniquely enriched in endocytosis (*q* = 4.09 x 10^−9^) and GABAergic synapses (*q* = 7.53 x 10^−3^), the cortical regions in components of the TCA cycle (*q* = 1.72 x 10^−3^) and the cerebellum in valine, leucine and isoleucine degradation pathways (*q* = 1.11 x 10^−6^) and fatty acids metabolism (*q* = 4.6 x 10^−4^) (Supplementary Table 7A-E). Together, these findings indicate that synapse proteome composition may reflect functional differences between interconnected brain regions.

### Regional differences in postsynaptic protein interaction networks

The organization and function of synapse proteomes have been studied using protein-protein interaction (PPI) networks^18,22,30,31^ and we used this approach to explore the organization of protein interaction networks in different brain regions. First, the total postsynaptic proteome network was built from the list of proteins obtained in this study and PPIs obtained by mining publicly available databases: BioGRID^32^, IntAct^33^ and DIP^34^ for both mouse and human. The total network consists of 1,016 proteins and 8,105 PPIs. Using the differential abundance of proteins as an edge weight, we constructed individual networks for each region, identified clusters and their corresponding enrichments in biological and disease functions (Materials and Methods, Supplementary Table 8). We found that each region-specific PPI network was split by the same method into a different number of clusters (cl. N), ranging from 82 to 112 clusters (results for a spectral clustering algorithm are shown in Supplementary Table 8). We assessed the clustering structure of each region’s PPIs for robustness and the resulting (consensus) clusters were examined for disease enrichment (Materials and Methods, Supplementary Table 9).

Examination of disease enrichment showed that some diseases impact clusters across all brain regions whereas others had more discrete regional effects. For example, all brain regions (except CxF) contained one highly significantly enriched cluster for autism spectrum disorder (adjusted *p*-values as follows): CB (cl. 19, *p* = 9.77 x 10^−8^), CxCA (cl. 9, *p* = 9.77 x 10^−8^), CxM (cl. 10, *p* = 2.76 x 10^−6^), Hip (cl. 53, *p* = 4.84 x 10^−7^), Hyp (cl. 48, *p* = 3.77 x 10^−6^) and ST (cl. 24, *p* = 8.17 x 10^−3^). We also found clusters associated with intellectual disability (ID) to be enriched in the CB (cl. 19, *p* = 4.36 x 10^−5^) and the ST (cl. 25, *p* = 6.64 x 10^−5^). We found that PPI clusters associated with bipolar disorder were moderately enriched in the CxF (cl. 50, *p* = 1.44 x 10^−2^) and Hyp (cl. 66, *p* = 2.05 x 10^−2^), while those associated with schizophrenia were highly enriched in the CB, CxCA and CxM (cl. 10, *p* = 1.29 x 10^−6^; cl. 9, *p* = 4.41 x 10^−6^; and cl. 10, *p* = 2.90 x 10^−5^, respectively) (see Supplementary Table 9 for all disease enrichment results).

We found regional variability in enrichment levels for neurodegenerative diseases. For Alzheimer’s disease, the most enriched cluster was in the Hip (cl. 53, *p* = 7.97 x 10^−6^), while for Parkinson’s disease a moderate enrichment occurred in the CxCA (cl. 32, *p* = 6.64 x 10^−3^). Clusters associated with dementia were significantly enriched only in the Hyp (cl. 6, *p* = 4.63 x 10^−5^).

### Identifying a stable core network

To define the network structures that are conserved across all brain regions, we identified 4,205 binary interactions (52% of total) that were found in the same cluster in the majority of regional networks (see Materials and Methods). We refer to this as the “stable” postsynaptic density (PSD) (1,016 proteins, Figure 5, Supplementary Table 10). Spectral clustering generated 73 clusters in total, where the nine largest represent crucial synaptic proteins and neural housekeeping functions: cl. 37 corresponds to the postsynaptic signaling complexes composed of MAGUK scaffold proteins, AMPA and NMDA receptors^23^, and other clusters contain ribosomal, metabolic enzymes and actin/myosin-associated proteins (Figure 5, Supplementary Table 10). We compared the composition of these stable communities with previously detected PPMs and found significant overlaps. For example, cl. 1 containing ATPase and cytochrome-related proteins and cl. 8 containing mitochondrial complex I proteins were over-represented in protein module PPM1 associated with related terms (*p* = 4.5 x 10^−4^ and *p* = 9.3 x 10^−5^, respectively); cl 4 is composed mainly of metabolic enzymes and falls almost entirely in to PPM1 associated with Parkinson’s disease, Alzheimer’s disease and metabolic pathways by KEGG pathway enrichment (Figure 5; *p* = 7.5 x 10^−5^). Molecules involved in memory and cognition (e.g. MAGUKs) that populated cl. 6 are distributed between PPM5 (*p* = 8.0 x 10^−3^) and PPM4 (*p* = 1.9 x 10^−2^) associated with the terms "neurotransmitter receptor binding and downstream transmission in the postsynaptic cell", "long-term potentiation" and "long-term depression".

**Figure 5.**
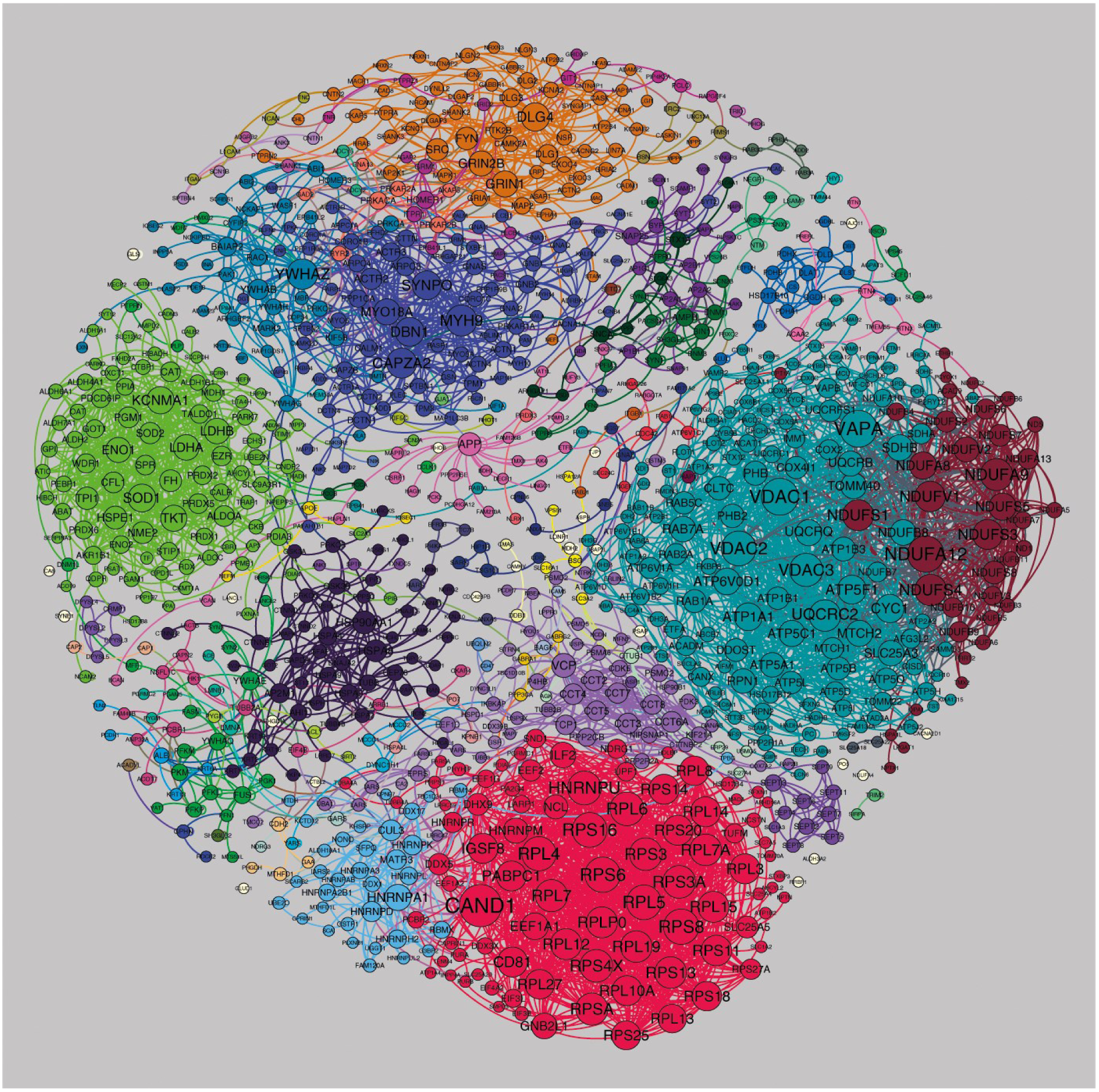
Postsynaptic proteome interaction network showing the cluster structure for the "stable network". A few large clusters with specific functionally related proteins could be detected: cl. 3 contains ribosomal proteins (red cluster at the bottom), cl. 4 contains metabolic enzymes (light green on the left), cl. 7 is enriched with actin-, myosin- and cytoskeleton remodeling-associated proteins (dark blue near the top), cl. 8 contains NADH-oxidoreductases (dark-red cluster on the right) and cl. 1 contains ATPases and voltage-dependent anion channels (light blue on the right). Cl. 6 corresponds to key proteins involved in synaptic transmission and plasticity, including AMPA, NMDA receptors and MAGUK proteins (orange cluster at the top). Networks were visualized with Gephi.

## Discussion

Using mass spectrometry we have examined the protein composition of the postsynaptic proteome of excitatory synapses from regions of the mouse brain and generated a freely available data resource (Edinburgh DataShare, http://dx.doi.org/10.7488/ds/1713). We found that a high percentage of proteins show abundant differences between brain regions. The postsynaptic proteome composition for each brain region forms a distinctive molecular signature. Because these proteomic data are obtained from tissue samples composed of many individual synapses, the proteomic signatures indicate that there might be synapse diversity at the single-synapse level. Consistent with this, we have recently examined the differential distribution of PSD95 and SAP102 in individual synapses across the whole mouse brain and found that these two proteins are differentially distributed into synapse subtypes^35^. Moreover, each brain region was composed of varying proportions of synapse subtypes, which results in each region having a “signature of subtypes”. Together, these findings indicate that postsynaptic proteome diversity seen at the level of brain regions arises at the individual synapse level.

The advantage of proteomic mass spectrometry is that it examines the expression of a large number of proteins and can therefore shed light on how sets of proteins are expressed. We found regional diversity in sets of proteins known to be associated with biochemical pathways controlling physiological processes (such as forms of synaptic plasticity and cognition) and diseases. We also found evidence that scaffold proteins involved with the supramolecular assembly of complexes and supercomplexes were differentially distributed across the brain.

The analysis of weighted PPI networks supports previous findings that the postsynaptic proteome has a modular structure^30^. We now find that the regional variability of protein complex composition strongly depends on the relative protein abundance, thereby providing the heterogeneity and unique biochemical signaling potential of each region. In each brain region we find that ~60% of the complexes/clusters are conserved in a stable network and that ~40% underpin regional specifications. Disease enrichment analysis, which was performed at the regional level, tends to identify the same clusters. Regional specificity results in effectively the same cluster being more or less enriched for each disease across the different brain regions.

The coordinate expression of sets and modules of proteins suggests the possibility that there is an underlying genetic mechanism coordinating the spatial expression of synapse proteins. Evidence in support of a coordinating genetic mechanism acting in the temporal domain was provided from transcriptome analyses of developing cultured neurons^36^ and lifespan expression data in the brain of mouse and human^2^, which show concerted regulation of postsynaptic proteins. Further evidence for an underlying genetic program regulating the differential spatial expression comes from the hierarchical clustering of protein expression that reveals a correlation with the early development of the nervous system. A similar result was obtained using the single-synapse resolution mapping: the regional signatures of synapse composition in the adult mouse brain are organized into three major groups corresponding to the earliest division of the neural tube. These multiple lines of evidence suggest that there is temporal and spatial regulation of postsynaptic proteome expression and that it produces diversity of synapse types^35^.

The purpose of regional diversity is most likely to subserve region-specific physiological and behavioral functions. It might also be to provide regional specializations within the greater systems-level organization or global circuitry of the brain. Our analysis of connectome data indicates that the anatomical circuitry of the brain links areas with distinct synapse proteome compositions. Given the differences in biochemical pathways, this indicates that functional specializations in regions are integrated by the connectome. This is consistent with our recent findings showing that the regional composition of the neocortex is linked to behavioral functions observed using functional magnetic resonance imaging (fMRI)^25^.

In conclusion, we provide a new data resource describing the composition of the postsynaptic proteome in excitatory synapses from regions of the mouse brain. These data indicate that molecular compositional differences in synapses in different brain regions are relevant to a broad range of physiological and disease processes.

## Acknowledgements

Support from the Medical Research Council (G0802238) and European Union Seventh Framework Programme (FP7 grant agreement no. 604102) and Horizon 2020 (grant agreement no. 72027). We thank T. Le Bihan and L. Imrie at SynthSys, University of Edinburgh, for mass spectrometry sample analysis. The LC-MS QExactive equipment was purchased by a Wellcome Trust Institutional Strategic Support Fund and a strategic award from the Wellcome Trust for the Centre for Immunity, Infection and Evolution (095831/Z/11/Z). Data were extracted from Neuroimaging Informatics Technology Initiative (NIFTI) files using a custom automated script written by Jeremy J. Roy, MEMEX, Inc. (Burlington, Ontario, Canada). We thank K. Elsegood for laboratory management and D. Maizels for artwork; C.S. Davey, editing.

## Author contributions

S.T.-G., J.D. brain samples; M.R. synapse biochemistry; M.R., O.S., C.M., J.D.A., bioinformatics; S.G.N.G. direction.

## Supplementary Materials

**Supplementary Figure 1.** The seven regions dissected from the mouse brain.

**Supplementary Figure 2.** The overlap between proteomes obtained in this study and previous studies.

**Supplementary Figure 3.** The principal component analysis (PCA) of the abundances of the 1,173 synaptic proteins quantified across the seven integral regions for all six mouse brains.

**Supplementary Figure 4.** Circos plots showing differential abundance of the six postsynaptic proteome modules (PPMs) across mouse brain regions.

**Supplementary Table 1.** LC-MS/MS quantitation and comparison of the mouse synaptic proteome across the seven integral regions of the mouse brain by one-way ANOVA.

**Supplementary Table 2.** New PSD proteins detected in this study compared to all other published mouse PSD studies.

**Supplementary Table 3.** Differential stability analysis and functional enrichment of synaptic proteins.

**Supplementary Table 4.** Summary of the 868 synaptic proteins found to be differentially abundant across the seven integral regions of the mouse brain.

**Supplementary Table 5.** Module gene list and abundances for all six modules.

**Supplementary Table 6.** Correlation of ABI functional connectivity with synaptic proteome abundances measured in this study.

**Supplementary Table 7.** Functional Enrichment of ABI and Roy Correlates by Brain region.

**Supplementary Table 8.** PSD protein interaction network analysis across the mouse brain.

**Supplementary Table 9.** Disease associated enrichment in the interactomes of the mouse PSD.

**Supplementary Table 10.** The “stable network”: postsynaptic proteome protein-protein interaction networks found to be enriched in at least four different regions of the mouse brain.

